# Embracing the dropouts in single-cell RNA-seq data

**DOI:** 10.1101/468025

**Authors:** Peng Qiu

## Abstract

One primary reason that makes the analysis of single-cell RNA-seq data challenging is dropouts, where the data only captures a small fraction of the transcriptome of each cell. Many computational algorithms developed for single-cell RNA-seq adopted gene selection and dimension reduction strategies to address the dropouts. Here, an opposite view is explored. Instead of treating dropouts as a problem to be fixed, we embrace it as a useful signal for defining cell types. We present an iterative co-occurrence clustering algorithm that works with binarized single-cell RNA-seq count data. Surprisingly, although all the quantitative information is removed after the data is binarized, co-occurrence clustering of the binarized data is able to effectively identify cell populations, as well as cell-type specific pathways. We demonstrate that the binary dropout patterns of the data provides not only overlapping but also complementary information compared to the quantitative gene expression counts in single-cell RNA-seq data.

## 1 Introduction

Single-cell RNA sequencing (scRNA-seq) is a powerful technology capable of unveiling cellular heterogeneity of the transcriptome at single-cell resolution, producing insights toward subpopulation structures and progression trajectories which would be hidden in bulk cell population RNA sequencing analyses [1, 2, 3]. Enabled by scRNA-seq advances including SMART-seq [4], CEL-seq [5], Dropseq [6], InDrop [7], Chromium 10X [8], SCI-seq [9] and SPLiT-seq [10], scRNA-seq is increasingly used and offers the promise of addressing a variety of biology questions, such as intra-population heterogeneity and novel subpopulation identification [11], developmental trajectories [12], and regulatory mechanisms [13, 14].

scRNA-seq experiments often generate large amounts of data, containing whole-genome gene expression measurements of thousands or more individual cells, which presents challenges in appropriate computational analysis and interpretation of the data [15]. There are several reasons why computational analysis of scRNA-seq data is challenging, such as high dimensionality, measurement noise, detection limit, unbalanced size between rare and abundant populations, etc. One important characteristic of scRNA-seq data that feeds into all these challenges is a phenomenon called “dropout”, where a gene is observed at a moderate expression level in one cell but is not detected in another cell of the same cell type [16]. These dropout events occur due to the low amounts of mRNA in individual cells and inefficient mRNA capture, as well as the stochasticity of mRNA expression. As a result of the dropouts, the scRNA-seq data is often highly sparse. The excessive zero counts cause the data to be zero-inflated, only capturing a small fraction of the transcriptome of each cell.

Many algorithms have been developed to analyze scRNA-seq data and computationally tackle the dropout problem. A common data preprocessing strategy is to focus on highly variable genes, apply principal component analysis (PCA) [17] for dimension reduction, apply t-Distributed Stochastic Neighbor Embedding (tSNE) [18] for visualization or further dimension reduction, followed by algorithms for clustering or trajectory identification [19, 20]. Examples include Seurat [6, 13, 21], MNN batch-effect-correction [22], scLVM [23], BackSPIN [24], STREAM [25], and many others. The choice of focusing on highly variable genes is based on the assumption that the cell-to-cell heterogeneity can be captured by genes exhibiting high variability, which is often true. However, the selection of highly variable genes can be sensitive to normalization and imputation, which affect the results of the subsequent clustering and trajectory analysis. In addition, genes that are not highly variable may also be useful for defining rare cell subpopulations.

There are also imputation algorithms designed to specifically address the dropouts. A few recent examples include CIDR, ZIFA, ZINB-WaVE, MAGIC, SAVER and scImpute. CIDR adjusts dropout values when computing pairwise distance among cells. The imputation is implicit because the adjustment of a particular gene in a cell changes each time when the cell is paired with a different cell [26]. ZIFA and ZINB-WaVE use zero-inflated models to achieve robust dimension reduction that accounts for the dropouts [27, 28]. MAGIC performs Markov diffusion to update one cell’s gene expression based on a weighted average of other cells. Such an update not only imputes the dropouts but also changes the non-dropout observations [29]. SAVER uses gene-to-gene relationships to impute the expression of each gene in each cell and provide uncertainty quantification for the estimated expression values [30]. scImpute uses cell-to-cell similarities and regularized regression to determine which zeros in the data are due to the dropout, and performs imputation only on the dropouts [31]. When applied to scRNA-seq data of heterogeneous cell populations, all these algorithms improved the separation among the subpopulations to certain extents.

In general, there are two schools of thoughts regarding the dropouts. One assumes that dropout affects the observed expression counts of all the genes in all the cells, such as MAGIC, which imputes all the zero and non-zero values in the data [29]. The other assumes that only a subset of the zeros are caused by dropout, such as scImpute, which identifies and imputes zeros that are due to the dropout [31]. It appears that MAGIC works better on data with underlying biological progression trajectories, while scImpute works better on data with cluster and subpopulation structures. However, there have been results supporting both ideas, and it remains unclear which one better describes the dropouts in scRNA-seq data.

Here, instead of treating the dropouts as a problem that needs to be fixed, we decide to embrace the dropouts as a useful signal. We first binarize the count matrix of the scRNA-seq data to be detected versus undetected, turning all the non-zero observations into one. All subsequent analyses only work with the binary version of the data. Therefore, the analysis pipeline does not require any normalization or imputation. We present an iterative co-occurrence clustering algorithm to cluster cells based on the binarized data. Although all the quantitative information of gene expression is completely removed after the data is binarized, we demonstrate that the co-occurrence clustering algorithm is able to effectively identify cell populations, as well as cell-type specific pathways beyond the highly variable genes. This suggests the binary dropout patterns in scRNA-seq data is as informative as the quantitative expression of highly variable genes.

## 2 Results

### Iterative co-occurrence clustering

The iterative co-occurrence clustering algorithm takes the binarized scRNA-seq count matrix as input and produces cell clustering results as output. A flowchart of the algorithm is shown in Figure 1. The algorithm works in a hierarchically divisive manner, and iteratively performs gene pathway identification and cell type discovery.

**Figure 1:**
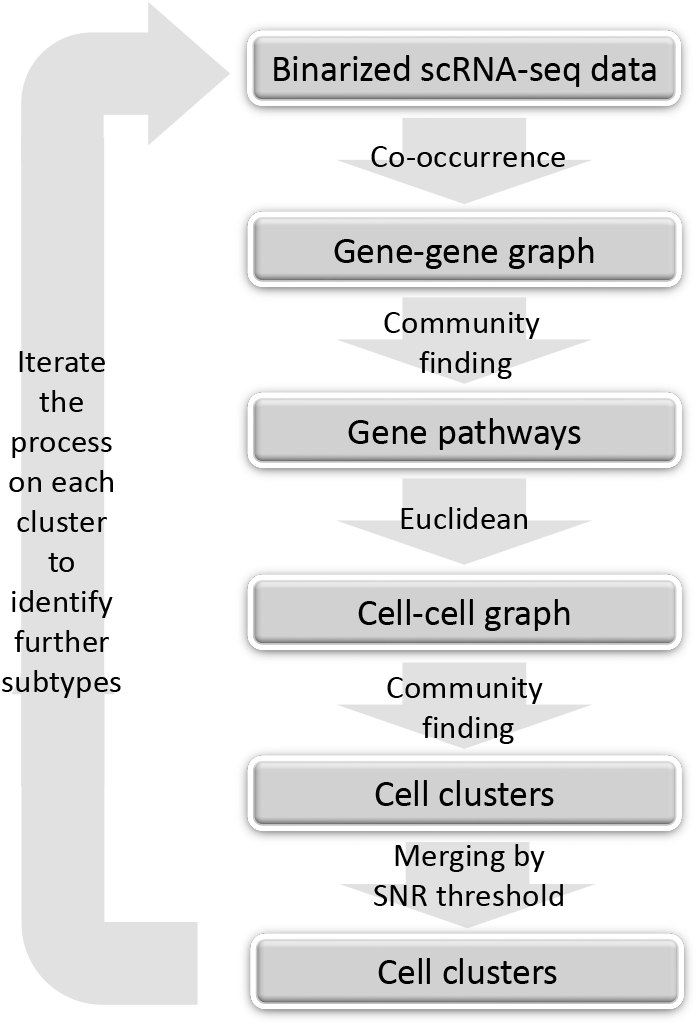
Flowchart of the iterative co-occurrence clustering algorithm.

The starting point of the algorithm is a root node at the top of a hierarchical tree, which contains all the cells in the data. The algorithm first evaluates a statistical measure for co-occurrence between each pair of genes, which is computed based on all the cells. The co-occurrence measures are filtered and adjusted by the Jaccard index [32], which defines a weighted gene-gene graph. The gene-gene graph is partitioned into gene clusters using community detection (e.g., Louvain algorithm [33]). The resulting gene clusters contain genes that share high co-occurrence based on all cells, and can serve as pathway signatures that separate major groups of cell types in the heterogeneous population. For each gene pathway, the percentage of detected genes is computed for each cell. These percentages form a low-dimensional representation of the cells, where each dimension describes the activities of one gene pathway in the cells. The algorithm then builds a cell-cell graph using Euclidean distances based on the low-dimensional pathway activity representation, uses the Jaccard index to filter the cell-cell graph, and applies community detection again to partition the cell-cell graph into cell clusters. For each pair of cell clusters, three metrics (mean difference, mean ratio and signal-to-noise ratio) are used to evaluate whether any of the gene pathways show differential activities. If none of the gene pathways exhibit different activity between the two clusters, these two cell clusters are merged. After merging the cell clusters according to pathway activities, each pair of the resulting cell clusters have at least one gene pathway that shows large difference between the two cell clusters. These cells clusters form children nodes of the root node, and are expected to capture the major groups of cell phenotypes in the data.

In the subsequent iterations, each resulting cell cluster (children node of the root node) can be further divided using the same process, which is expected to identify smaller subtypes in each major group of cell phenotypes. This is because when the gene-gene co-occurrence graph is computed based on only the cells in a children node, the significant edges are driven by genes that separate the major cell subtypes inside this children node. These subtypes form lower-level children nodes further down the hierarchical tree, which are examined in later iterations of the algorithm. If a children node is not further divided because the community detection step produces only one cell cluster or all cell clusters are merged, this children node becomes a leaf of the hierarchial clustering process, and is reported as one cell type identified by the algorithm. Therefore, the merging criteria define when the iterations stop and the final number of clusters identified by the algorithm. More details are described in Methods.

Similar to all divisive hierarchical clustering methods, the co-occurrence clustering algorithm represents a divide-and-concur strategy. When examining the root node that contains many cell types, the gene-gene co-occurrence measures and the gene pathways are dominated by the differences among the major groups of cell types, enabling the initial iteration to identify the major cell clusters. Each subsequent iteration focuses on a previously identified cell cluster, where the gene-gene co-occurrence measures lead to different gene pathways driven by the heterogeneity within the cell cluster, which provides the basis for identifying further cell cluster. The key strength of this algorithm is that different iterations use different sets of genes and pathways to cluster the cells.

### Co-occurrence clustering applied to PBMC

To evaluate its utility in analyzing scRNA-seq data, the co-occurrence clustering algorithm was applied to a dataset of Peripheral Blood Mononuclear Cells (PBMC) freely available from 10X Genomics (https://s3-us-west-2.amazonaws.com/10x.files/samples/cell/pbmc3k/pbmc3k_filtered_gene_bc_matrices.tar.gz). This dataset contains scRNA-seq counts data for 32,738 genes in 2,700 single cells that were sequenced on the Illumina NextSeq 500. 97.41% of the count matrix were zeros.

The first iteration examined the initial cluster 0 which contained all the 2,700 cells. The algorithm constructed a gene-gene graph based on co-occurrence. Community detection in the gene-gene graph led to four gene pathways, indicated by the upper-right heatmap in Figure 2(a). Each pathway contained genes that were significantly co-detected, as shown in the upper-left heatmap of Figure 2(a). The number of genes in the four pathways ranged from 106 to 538. For each gene pathway, the percentage of detected genes was used to represent the detected activity of the pathway in individual cells, which was shown in the bottom heatmap of Figure 2(a). Based on the cell-cell graph constructed by the Euclidean distance of the pathway activity representation, community detection analysis yielded fifteen cell clusters, which were subsequently merged to four cell clusters according to the merging thresholds of 1.5, 0.5, 2 for signal-to-noise ratio, mean difference and mean ratio (see details in Methods). The upper-left heatmap in Figure 2(a) showed the binarized count data of the genes in the pathways across all the individual cells, where the rows and columns were arranged by the gene pathways and cell clusters identified by the algorithm. It was obvious that distinct cell clusters can be defined by the binary data of the genes in the identified pathways, both from the heatmap of the binarized data itself and the heatmap of the pathway activity percentages.

**Figure 2:**
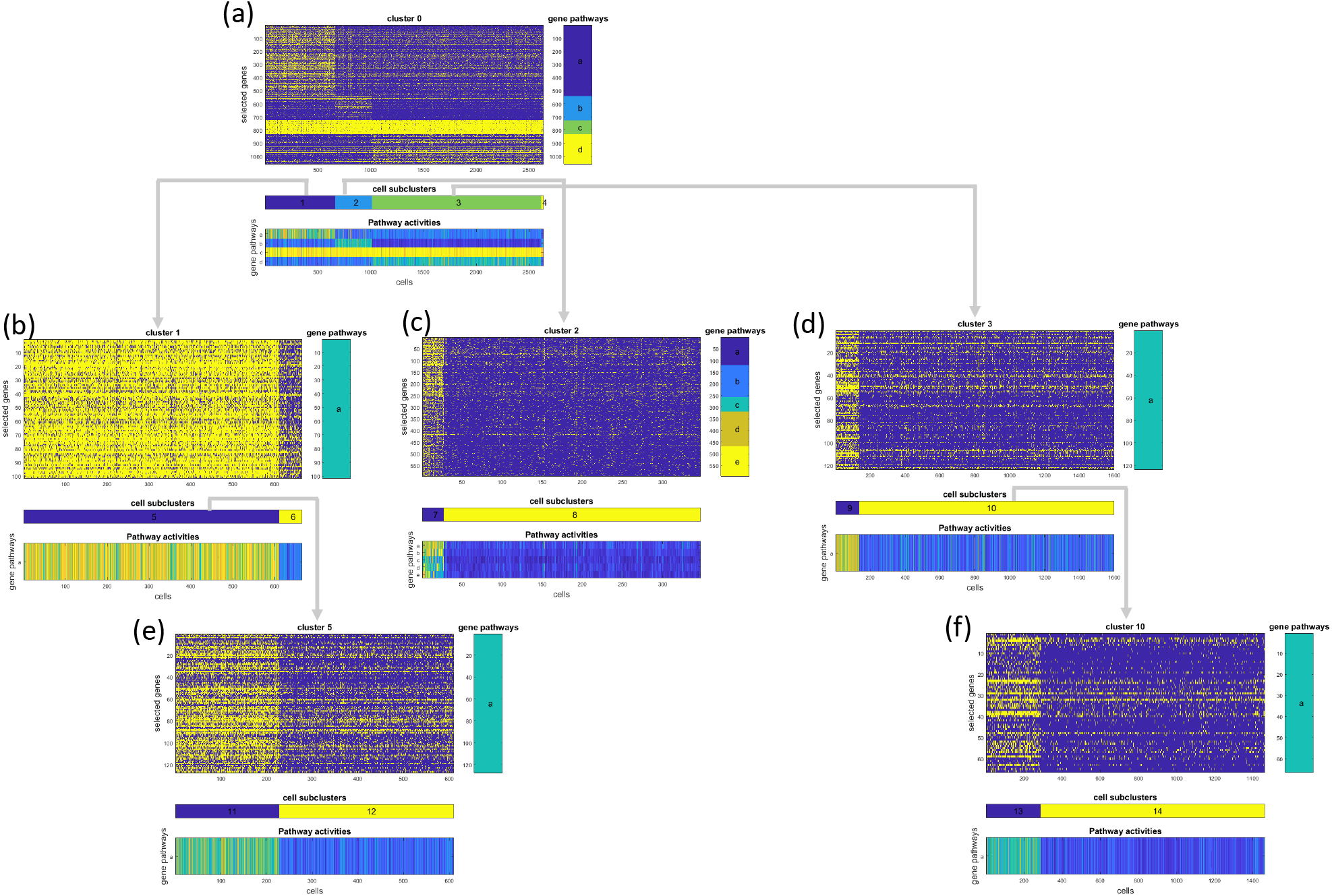
Co-occurrence clustering applied to binarized PBMC data generated by 10X.

In the following iterations, the four cell clusters identified in the first iteration were separately examined by the same algorithm. Cell clusters 1, 2 and 3 were further divided into smaller clusters, as shown in Figure 2(b-d). In each case, the identified gene pathways contained fewer genes compared to the first iteration. This was because the first iteration examined all cells which contained highly diverse cell types with large differences reflected in many genes, whereas the cell clusters examined in the following iterations were less heterogeneous and the differences among further subtypes were more subtle. Clusters 4 was not divided because the algorithm did not identity any gene pathway with genes that exhibited significant co-occurrence.

Moving down the hierarchical process of the iterative algorithm, among cell clusters 5 ~ 10, only clusters 5 and 10 were further divided, as shown in Figure 2 (e-f). The others were not divided because of either lack of gene pathways identified in the gene-gene graph, lack of cell clusters identified in the cell-cell graph, or lack of further clusters that exhibit differential pathway activities exceeding the thresholds in the merging step. Overall, the co-occurrence clustering algorithm identified 9 cell clusters using the binarized PBMC dataset.

This dataset has been previously analyzed with Seurat, which identified 8 cell clusters using community detection based on principle component analysis of the expression data of highly variable genes, https://satijalab.org/seurat/pbmc3k_tutorial.html. The clustering results between cooccurrence clustering and Seurat were highly similar, with a Rand Index of 0.85. A detailed comparison of the two clustering results was visualized in the heatmap in Figure 3(a). Each column of the heatmap was colored by the percentage overlap between one co-occurrence cluster and the Seurat clusters. Most of the columns were dominated by a single entry, indicating that each co-occurrence clusters was primarily enriched by one Seurat cluster. Five Seurat clusters (Dendritic cells, CD4 T cells, CD8 T cells, NK cells, FCGR3A+ Monocytes) were captured by individual co-occurrence clusters (7, 14, 13, 9, 11). Two Seurat clusters (B Cells, CD14+ Monocytes) were divided into subtypes. Overall, except for the Megakaryocytes, most of the Seurat clusters were well-separated in the co-occurrence clustering results. Since both algorithms had tuning parameters that affected the number of resulting clusters, the difference in the number of clusters was not indicative. However,the general agreement between the two clustering results was striking, and suggested that in terms of defining cell types, the binary dropout patterns of the data was as informative as the quantitative expression of highly variable genes.

**Figure 3:**
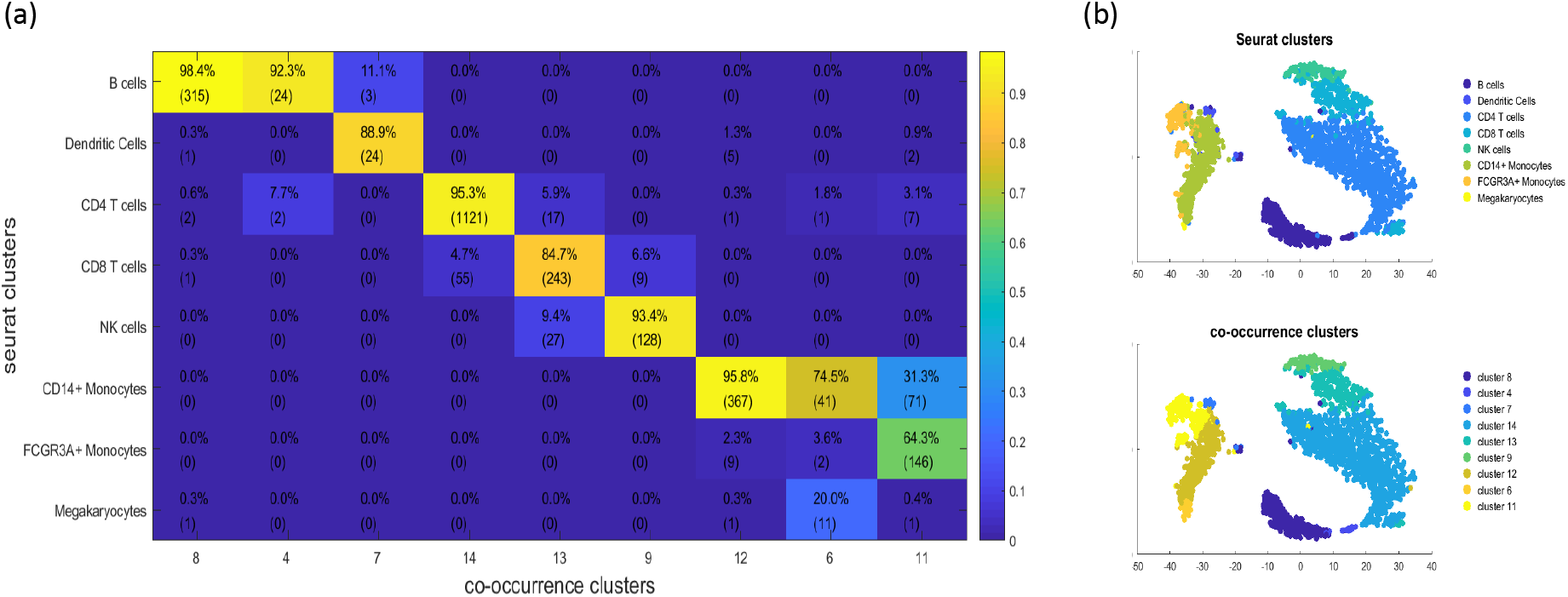
Comparison between co-occurrence clusters and Seurat clusters on PBMC data. (a) In the confusion heatmap, each column was colored by the percentages of cells in one co-occurrence cluster that belonged to each of the Seurat clusters. The integer values in parentheses represented the number of cells. (b) tSNE was applied to the percentage detection of gene pathways, embedding the cells according to the pathway activity space. Cells were colored by the clustering results from Seurat and co-occurrence clustering.

In the analyses of this dataset, the number of highly variable genes defined by Seurat was 1838, and the 13 gene pathways identified by co-occurrence clustering contained a total of 1546 genes. The overlap was 359. Many highly variable genes were not included in the co-occurrence gene pathways mainly because co-occurrence clustering used binarized data. For the genes whose expression were quantitative informative in defining cell types, their binary dropout patterns might not be informative. In contrast, co-occurrence clustering utilized many genes that were not highly variable.

Using the 13 gene pathways defined by co-occurrence clustering, the percentages of detected genes of the gene pathways were calculated for each cell, which led to a representation of cells in the pathway activity space. In Figure 3(b), tSNE was applied to visualize the cells in the pathway activity space, with the clustering results overlaid by color. In the tSNE visualization, the major cell types were separated, which again showed that the binarized data contained information for delineating the major cell types in PBMC.

In addition to the percentages of detection, the activities of gene pathways can also be quantified by the average expression level of the detected genes. In Figure 4, these two metrics were compared for all the 13 co-occurrence gene pathways identified from this PBMC dataset. The five gene pathways in the top row of Figure 4 showed strong positive correlations, meaning that the percentage of detection and the detected expression level of these five pathways provided similar information. However, for the remaining pathways, percentage of detection and average expression level were less correlated. In the middle row of Figure 4, the colors showed that some of the co-occurrence cell clusters were separated according to percentage of detection, but had overlapping expression levels. This observation suggested that the percentage of detection provided additional information complementary to the detected gene expression level.

**Figure 4:**
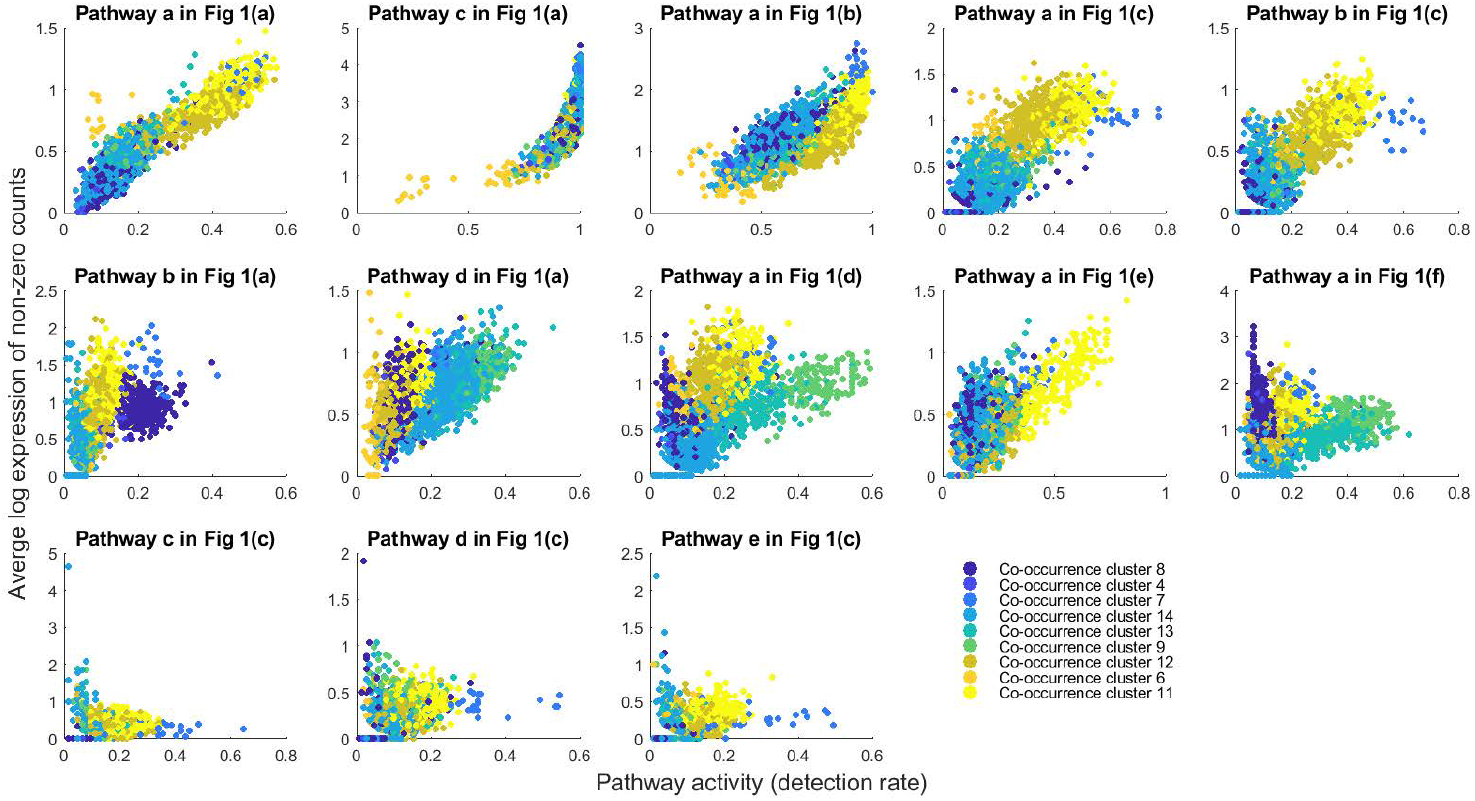
Relationship between percentage of detection and quantitative gene expression. Each panel visualizes the cells according to one co-occurrence gene pathway. The horizontal axis is the percentage of genes detected in the pathway, and the vertical axis is the average log-transformed counts for the detected genes in the pathway. Cells are colored by the co-occurrence clusters.

### Co-occurrence clustering applied to mouse inner ear sensory epithelia

The co-occurrence clustering algorithm was further tested on an scRNA-seq dataset on the cellular complexity in the mouse inner ear sensory epithelia, which contained hair cells (HCs) and supporting cells (SCs) organized in exquisite mosaic patterns to perform sensory functions, transitional epithelial cells (TECs) located at the border between sensory and non-sensory regions, as well as non-sensory cells (NSCs) [34]. The data was generated with SMART-seq2, and provided gene expression counts for 26,585 genes across 321 single cells, where the percentage of zero counts was 77%.

Figure 5 showed the iterative process of co-occurrence clustering applied to this dataset. Each iteration identified 1 ~ 5 gene pathways that contained significantly co-detected genes with respect to the set of cells under consideration. The bottom heatmap for each iteration showed clear distinction among the cell clusters identified in the iteration, and hence, provided strong evidence to support the co-occurrence clusters. Figure 6(a) showed the comparison between the co-occurrence clusters and the experimentally defined cell types in this dataset. Co-occurrence clustering produced a total of 10 clusters, which separated the four experimentally defined cell types (HC, SC, TEC and NSC) and identified potential cell subtypes. For example, the SCs were divided into five co-occurrence clusters (5, 8, 9, 11, 15), which were well-separated in the tSNE visualization of the pathway activity space shown in Figure 6(b).

**Figure 5:**
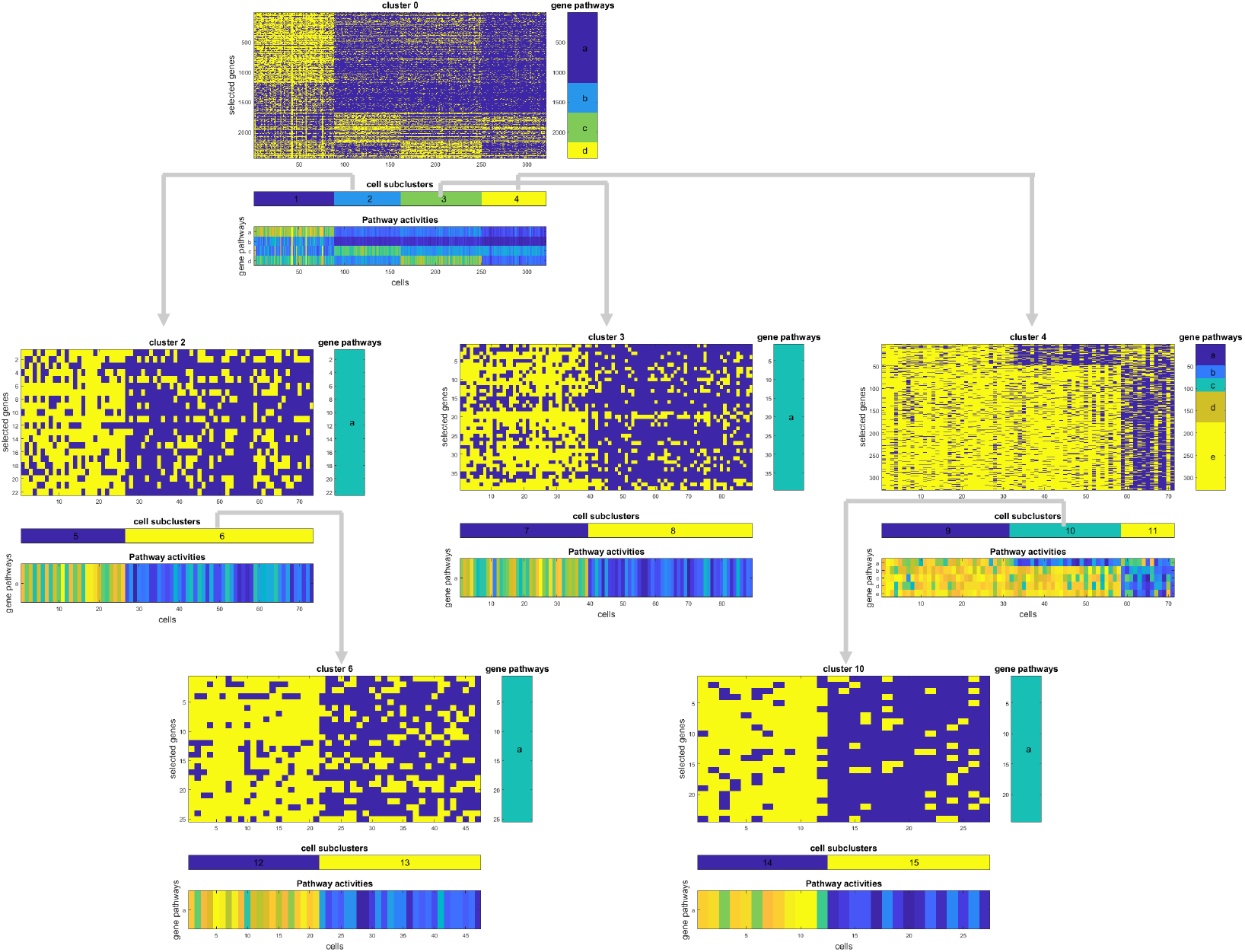
Co-occurrence clustering applied to binarized scRNA-seq data of mouse inner ear sensory epithelia.

**Figure 6:**
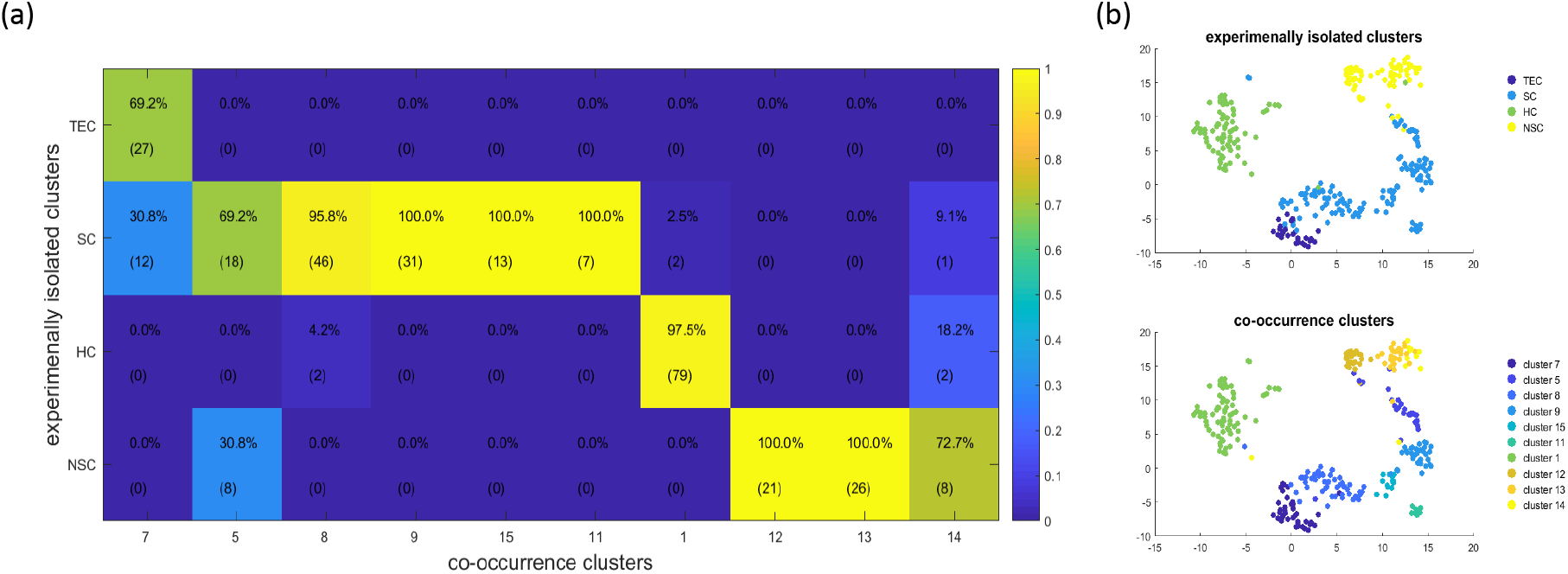
Comparison between co-occurrence clusters and experimentally isolated cell types in the dataset of mouse inner ear sensory epithelia. (a) In the confusion heatmap, each column was colored by the percentages of cells in one co-occurrence cluster that belonged to each of the experimentally defined cell type. The integer values in parentheses represented the number of cells. (b) tSNE was applied to the percentage detection of gene pathways, embedding the cells according to the pathway activity space. Cells were colored by the experimental clusters and the co-occurrence clusters.

Similar to the previous analysis of the PBMC dataset, the gene pathways in the initial iteration contained relatively large numbers of genes, and the size of gene pathways decreased as the algorithm iterated to lower levels of the hierarchical process. This was because the initial iteration examined all cells in the dataset, which contained drastically distinct cell types that can be characterized by the binary on/off expression states of many genes. The subsequent iterations examined cell clusters derived from previous iterations, where the remaining heterogeneity typically manifested in relatively smaller number of genes.

### Co-occurrence clustering applied to the human prefrontal cortex

Co-occurrence clustering was applied to an scRNA-seq dataset of the developing human prefrontal cortex, generated with SMART-seq2 [35]. This dataset was previously analyzed by a combination of tSNE and Seurat, which defined six major clusters: neural progenitor cells (NPCs), excitatory neurons, interneurons, astrocytes, oligodendrocyte progenitor cells (OPCs) and microglia [35]. The data consisted of measurements for 24,153 genes across 2,394 single cells, and the dropout rate was 82%. When applied to this dataset, the co-occurrence clustering algorithm went through 25 iterations with meaningful gene pathways and cell clusters, and eventually identified a total of 35 cell clusters. The visualization of the co-occurrence clustering process contained five times more plots compared to Figure 2, which were available in the supplementary materials. A comparison of the Seurat clusters and co-occurrence clusters is shown in Figure 7. Again, each column was dominated by one entry, which showed that each co-occurrence cluster was nested within one Seurat cluster. Several rows contained multiple entries close to 100%, indicating that the co-occurrence algorithm divided the Seurat clusters into smaller clusters.

**Figure 7:**
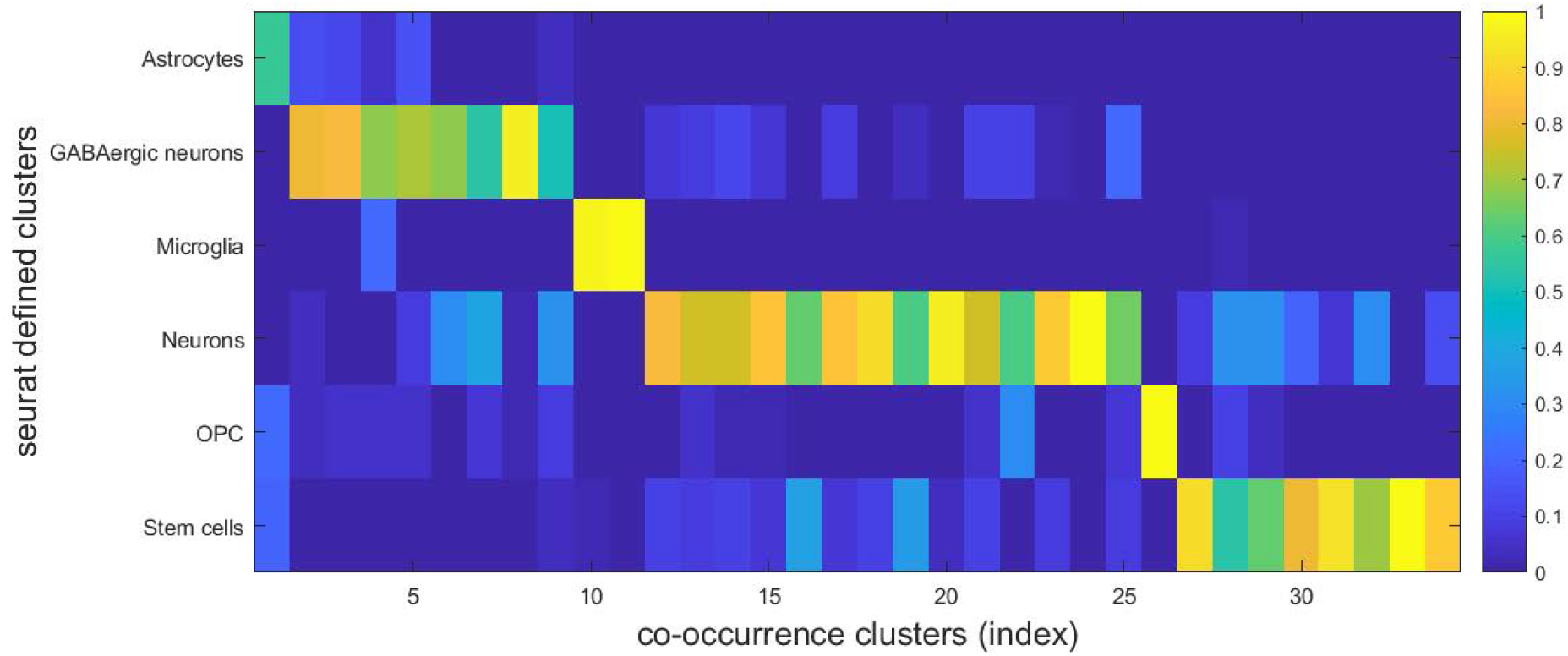
Comparison between co-occurrence clusters and seurat clusters in the human prefrontal cortex dataset. Each column was colored by the percentages of cells in one co-occurrence cluster that belonged to each of the experimentally defined cell type.

### Co-occurrence clustering applied to separate mouse tissue types

To demonstrate the utility and scalability, the co-occurrence clustering algorithm was applied to the Tabula Muris, which contained scRNA-seq data for about 100,000 cells from 20 organs and tissues in mouse [36]. The organs included are skin, fat, mammary gland, heart, bladder, brain, thymus, spleen, kidney, limb muscle, tongue, marrow, trachea, pancreas, lung, large intestine, and liver. Many of these organs were processed using two methods, SMART-seq2 on FACS-sorted cells and microfluidic droplets from 10X Genomics. The FACS-sorted SMART-seq2 dataset contained count data for 23,433 genes across 53,760 cells, with an overall dropout rate of 89%. The droplet-based 10X dataset contained count data of 70,118 cells for the same 23,433 genes, with an overall dropout rate of 93%. The Tabula Muris allowed evaluation of co-occurrence clustering on datasets with similar underlying heterogeneity but profiled by two different scRNA-seq technologies.

Co-occurrence clustering was applied to the droplet-based dataset and the FACS-based dataset separately. In both datasets, co-occurrence clustering identified more than 100 cell clusters. Visualizations of each iteration of the co-occurrence clustering processes were available in the supplementary materials. The Tabula Muris provided tissue type annotations for each individual cell, which was used to evaluate whether the co-occurrence clustering algorithm was able to delineate various tissue types. As shown in Figure 8, in both datasets, co-occurrence clustering successfully separated the tissue types in both datasets, and identified further subpopulations within many of the tissue types. Figure 9 compared the numbers of subpopulations co-occurrence clustering identified within each of the overlapping tissue types in the two datasets, which were generally in line with each other for many of the tissue types. However, there were also marked differences. For example, co-occurrence clustering identified multiple cell clusters within trachea, lung and spleen in the droplet-based dataset, but only one cell cluster for each in the FACS-based dataset. This was partially because the distributions of cells across the tissue types were different between the two datasets. Trachea, lung and spleen were the dominant populations that accounted for 30%, 13% and 14% of the droplet-based dataset, whereas these three tissue types together summed up to 9% of the cells in the FACS-based dataset. Despite the differences in the number of identified subpopulations between the two dataset, this analysis demonstrated that co-occurrence was able to work with large scRNA-seq datasets generated by different technologies, and identify tissue types and subpopulations beyond the tissue types based on the binary dropout patterns in the data.

**Figure 8:**
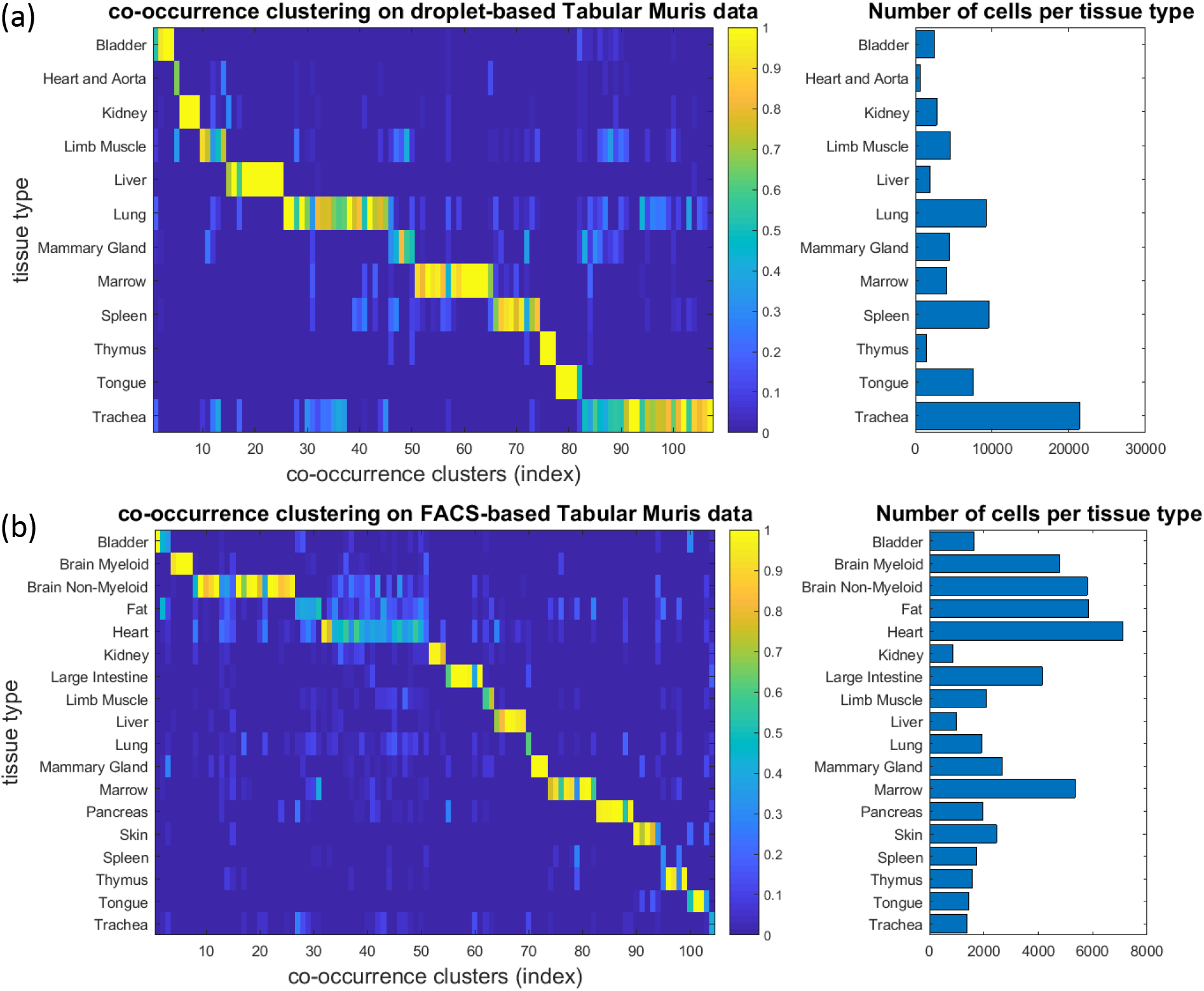
Co-occurrence clustering applied to binarized scRNA-seq data of the Tabula Muris. (a) Cooccurrence clustering separated tissue types in the droplet-based scRNA-seq data. (b) Co-occurrence clustering separated tissue types in the FACS-based scRNA-seq data.

**Figure 9:**
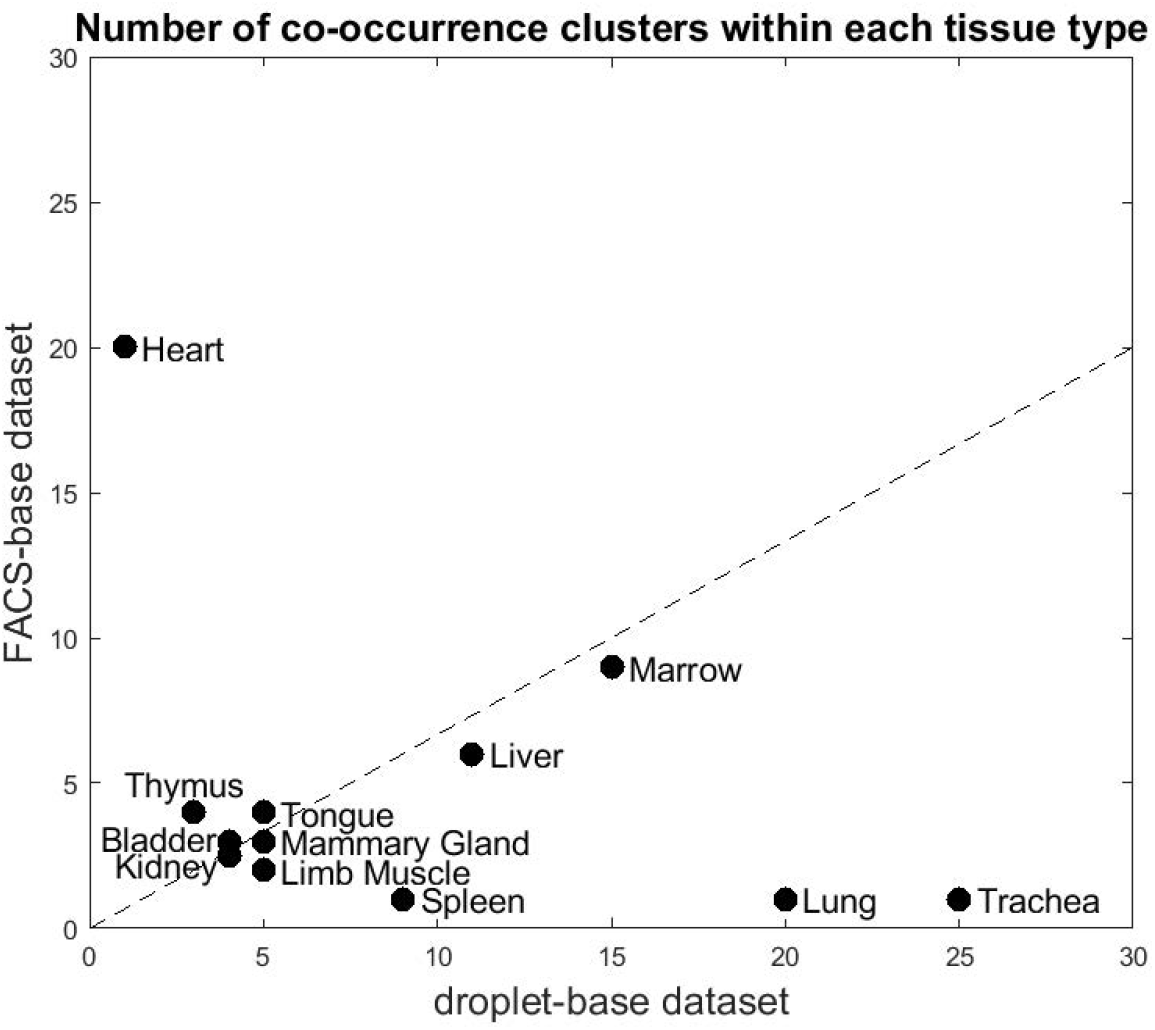
Numbers of co-occurrence clusters identified within the overlapping tissue types between the droplet-based and FACS-based dataset in the Tabula Muris.

## 3 Discussion

In this paper, the co-occurrence clustering algorithm was developed to identify cell clusters based on binarized scRNA-seq data. The binarized data represented the dropout patterns in the data, namely which genes were detected or undetected in each individual cell, while the quantitative gene expression information was removed. Most existing computational methods for scRNA-seq treated dropout as a problem to be fixed. Here, an opposite view was explored. The co-occurrence clustering algorithm embraced dropout as a useful signal for defining cell types. Using several scRNA-seq datasets generated by different scRNA-seq technologies in various biology contexts, we demonstrated that co-occurrence clustering on binarized data was able to effectively identify cell clusters, which largely agreed with clusters derived from computational analysis of highly variable genes or experimentally defined tissue types. These results provided strong evidences that the binary dropout patterns in the data is a useful signal, and is as informative as the quantitative expression of highly variable genes.

In existing scRNA-seq analysis methods, feature selection is typically performed only once, as a form of data preprocessing before running subsequent clustering algorithms. The two most popular feature selection strategies are using highly variable genes and performing principal component analysis, both of which are primarily driven by genes that exhibit high expression and/or high variation. In the co-occurrence clustering algorithm, feature selection is re-visited in each iteration. Depending on the set of cells under consideration, each iteration builds a gene-gene graph based on co-occurrence, and applies community detection to identify gene pathways that are able to characterize the heterogeneity in the set of cells under consideration. As discussed in the Results section, the gene pathways constructed in the initial iteration are often of larger size compared to those in later iterations. This is mainly because the initial set of cells is typically more heterogeneous than the sets of cells considered in later iterations. This observation suggests that different gene sets or pathways are optimal for analyzing cell populations with different amount of heterogeneity.

In almost all of the datasets analyzed here, the co-occurrence clustering algorithm generated more clusters than the previous analyses of these datasets, which can be both a blessing and a curse. Although the dropout patterns showed clear differences among the clusters generated in each iteration (Figures 2, 5, and supplementary materials), the results presented a challenge for interpreting the biological functions and distinctions among the cell clusters. One strategy to reduce the number of cell clusters is to bring back the quantitative gene expression information as a post-processing analysis, and merge cell clusters that do not have significantly differentially expressed genes. Another strategy is to put constraints on the gene pathways, such as removing genes associated to unwanted variations among cells (e.g. cell cycle), requiring gene pathways to be sufficiently large, or prescribing gene pathways by incorporating prior knowledge, which are all possible directions for extending the co-occurrence clustering algorithm.

## 4 Methods

### Datasets

The Peripheral Blood Mononuclear Cells (PBMC) dataset was downloaded from 10X Genomics (https://s3-us-west-2.amazonaws.com/10x.files/samples/cell/pbmc3k/pbmc3k_filtered_gene_bc_matrices.tar.gz). The datasets on mouse inner ear sensory epithelia and human prefrontal cortex were both downloaded from GEO, with accession numbers GSE71982 and GSE104276, respectively. The Tabula Muris was obtained from the easy-data Github repository provided in the collaborative computational tools for the Human Cell Atlas (https://github.com/czi-hca-comp-tools/easy-data/blob/master/datasets/tabula_muris.md).

### Convert scRNA-seq count matrix into binary

The only data preprocessing required here is converting the count matrix into binary, where all the dropouts are still 0, and all the non-zero counts are turned into 1 regardless of the expression level. No normalization, transformation or imputation is required.

### Filter genes and cells before co-occurrence clustering

In each iteration, the co-occurrence clustering algorithm focuses on the binarized expression data of one cell cluster. Based on the binary data, the algorithm filters both genes and cells. More specifically, the algorithm removes genes detected in too few cells, and removes cells in which too few genes are detected. The default thresholds for both genes and cells were 10 for all the datasets analyzed in this paper.

### Construct gene-gene graph and gene pathways

Given the set of cells under consideration in the current iteration, the co-occurrence clustering algorithm first evaluates the co-occurrence between each pair of genes using the chi-square statistics. Based on the binarized data of two genes *g*_1_ and *g*_2_, denote *A* as the number of cells in which both genes are detected, *B* as the number of cells in which *g*_1_ is detected but *g*_2_ is undetected, *C* as the numbers of cells where *g*_1_ is undetected but *g*_2_ is detected, and *D* as the number of cells where both genes are undetected. The chi-square score is defined as 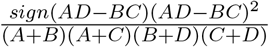. If the score is large, the two genes exhibit high co-occurrence. Random permutation is used to the define a threshold for what score is considered as large. 10 randomly permutated datasets are obtained by randomly reshuffling each row/gene independently, and the highest chi-square score from the random data is recorded. In the collection of pairwise chi-square scores from the actual data, for the ones smaller than the best random score, the mean and standard deviation are computed. A threshold is defined as the mean plus the standard deviation. An undirected unweighted gene-gene graph is then constructed by applying the threshold to the pair-wise chi-square scores. The Jaccard index [32] is applied to filter the unweighted graph into a weighted graph, and the Louvain algorithm [33] is applied to detect communities in the gene-gene graph, which are referred to as gene pathways here. A pre-defined threshold is used to discard gene pathways that are too small. If all the gene pathways produced by the community dection algorithm are smaller the threshold, all of them are discarded, and the current iteration ends without dividing the cells under consideration into clusters. The default threshold for the minimum pathway size was 20 for all the datasets analyzed in this paper.

### Construct cell-cell graph and cell clusters

For each gene pathway generated from the gene-gene graph, the percentage of detected genes is computed for each cell. These percentages form a low-dimensional representation of the cells, where the dimensionality is the number of gene pathways, and each dimension describes the activity of one gene pathway in the cells. Using the pairwise Euclidean distance among the cells based on the pathway activity representation, a k-nearest neighbor graph is constructed, which is an undirected unweighted cell-cell graph. The Jaccard index is again applied to filter the unweighted graph into a weighted graph, and the Louvain algorithm is applied to detect communities in the cell-cell graph, which are referred to as cell clusters here. Cell clusters that are smaller than a pre-defined threshold are considered as tiny clusters, and are merged into the nearest non-tiny cluster. Here, “near” is defined by Euclidean distance based on the pathway activity representation. If all cell clusters are tiny, or only one cluster remains after the tiny clusters are merged, the current iteration ends without generating new clusters. In all the datasets analyzed in this paper, the default k was 5 for the k-nearest neighbor graph, and the default threshold for tiny clusters was 10.

### Merge cell clusters

Although the cell-cell graph is defined based on the pathway percentages of detection, the cell clusters generated by community detection on the cell-cell graph are not necessarily prominently different in terms of the pathway percentages of detection. Therefore, the algorithm further merges the cell clusters according to three metrics of the percentages of detection: mean difference, mean ratio, signal-to-noisy ratio (SNR). For two cell clusters, the mean difference of a gene pathway is defined as the difference in the mean of the pathway’s percentage of detection in the two cell clusters; the mean ratio of the gene pathway is the ratio between the mean of the pathway’s percentage of detection in the two cell clusters; the SNR of the pathway is defined as the mean difference over the sum of the standard deviations of pathway’s the percentage of detection in the two cell clusters. In order for two cluster to be considered as prominently different, the algorithm requires two criteria to be met: (1) the maximum of their SNRs of the gene pathways is larger than 1.5, and (2) either the maximum of their mean differences for the gene pathways is larger than 0.5, or the maximum of their mean ratios is larger than 2. Clusters that do not meet these criteria are merged. After the clusters are merged according to these criteria, any two resulting cell clusters will exhibit prominent difference in at least one gene pathway, where prominent difference means that the SNR is larger than 1.5, and either the mean difference is larger than 0.5 or the mean ratio is larger than 2. If only one cluster remains due to these merging criteria, the current iteration ends without generating new clusters. If otherwise, subsequent iterations of the algorithm will examine the resulting cell clusters separately, and see whether they can be further divided in to sub-clusters that are prominently different in terms of certain gene pathways. The values 1.5, 0.5 and 2 were the default for all the datasets analyzed in this paper.

## Availability

Source code for the co-occurrence clustering algorithm implementation is available at https://github.com/pqiu/cooccurrence_clustering.

## Acknowledgement

The author gratefully acknowledges funding from the Chan Zuckerberg Initiative and the National Science Foundation (CCF1552784). P.Q. is an ISAC Marylou Ingram Scholar.

